# Aphid stylin cuticular proteins contribute to turnip mosaic virus (*Potyvirus*) transmission

**DOI:** 10.64898/2026.03.13.711641

**Authors:** Yu Fu, Emma Achard, Baptiste Monsion, François Hoh, Sophie Le Blaye, Bastien Cayrol, Nicolas Sauvion, Gaël Thébaud, Gaël Le Trionnaire, Fangfang Li, Stefano Colella, Marilyne Uzest

**Author notes:** Corresponding authors (SC); (MU).

## Abstract

Hundreds of plant viruses are transmitted by aphid vectors, among which non-circulative ones are acquired and inoculated from one host to another within seconds. These viruses are retained on receptors located at the surface of the cuticle of aphid mouthparts. Members of the *Potyviridae* family are the most abundant RNA viruses infecting plants, and they cause significant economic losses. Among them, viruses of the *Potyvirus* genus are transmitted by aphids in a stylet-borne manner. Their receptors in aphid stylets remain poorly characterized. Using turnip mosaic virus (TuMV, *Potyvirus rapae*) as a model, we developed complementary approaches to investigate potyvirus-aphid interactions in three vector species. Immunofluorescence detection and transmission electron microscopy revealed the presence of TuMV in the distal part of aphid maxillary stylets, both in the food canal and on the acrostyle. This cuticular organ houses Stylin proteins, including Stylin-01, the receptor for the cauliflower mosaic virus (*Caulimovirus tesselobrassicae*). Using CRISPR-Cas9-edited Stylin-01 mutant lines in the pea aphid, we demonstrated that this protein plays an important role in TuMV transmission. Complementary RNA interference silencing experiments revealed that Stylin-04/04bis also mediate TuMV transmission. Furthermore, our findings reveal that targeting simultaneously Stylin-01 and Stylin-04/04bis more strongly impaired the aphids’ ability to transmit TuMV, suggesting that virus transmission relies on a multi-component stylin interface rather than a single receptor. In conclusion, these results highlight that in complex interactions between potyviruses and their aphid vectors, Stylin proteins are key actors, underscoring their importance in the transmission of stylet-borne viruses.

**Author summary:** Potyviruses are the largest group of plant-infecting RNA viruses and major threats to crops worldwide. Aphids can transmit them within seconds retaining virus particles on their needle-like mouthparts through specific interactions at the stylet surface. However, the identity of the aphid proteins involved in this process has remained unknown. Using turnip mosaic virus (TuMV), a potyvirus with an exceptionnally broad host ranges, we provide evidence of virus retention at the acrostyle, a specialized cuticular microstructure at the tip of aphid stylets. CRISPR-Cas9 stable aphid mutant lines and RNAi-mediated gene silencing revealed that the acrostyle cuticular proteins Stylin-01 and Stylin04/04bis promote TuMV transmission. These findings uncover previously unknown molecular components of the aphid-potyvirus interface and identify potential targets to disrupt virus-vector interaction and virus transmission.

## Introduction

Potyviruses constitute the largest group of plant-infecting RNA viruses. They are among the most economically damaging plant viruses worldwide, impacting host plant physiology and causing substantial losses across a wide range of crops [1]. Aphids transmit them in a non-circulative, non-persistent manner [2]. Potyviruses can bind to aphid stylets and are acquired or released quickly within seconds to a few minutes. Early experimental studies showed that chemical or ultraviolet treatments applied to the distal tip of aphid stylets abolished transmission of potato virus Y (PVY, *Potyvirus yituberosi*, *Potyviridae*), suggesting that the virus is retained at or near the stylet tips [3,4]. These initial observations were later confirmed through transmission electron microscopy (TEM) and immunolabeling approaches, which revealed the presence of tobacco etch virus (TEV, *Potyvirus nicotianainsculpentis*) and PVY at the distal tip of the stylets of their aphid vector *Myzus persicae* [5–7]. Potyviruses interact with the stylet of aphids according to the helper strategy, where the viral protein helper component proteinase (HC-Pro) establishes a reversible “molecular bridge” between the viral capsid and receptors present in the stylets [8]. HC-Pro is a 50 kDa multifunctional protein consisting of three distinct domains: the helical-rich N-terminal domain (amino acids 1-100) is notably involved in aphid-mediated transmission [9]; the central domain (amino acids 101-299) participates in RNA silencing suppression, genome amplification, symptom severity, and viral movement [10,11]; and the helical-rich C-terminal domain (amino acids 300-458), which exhibits proteolytic activity, interacts with the viral capsid [11–14]. Specifically, two conserved motifs of HC-Pro are paramount for the bridging function: KITC likely interacts with receptors in aphid stylets, and PTK interacts with DAG present in the coat protein [9,15,16]. Oligomers of HC-Pro form via cysteine bridges [14,17–19] and are associated with active transmission of potyviruses by aphid vectors [17,18].

Despite the central role of vector transmission in the potyvirus life cycle, no specific receptor for potyviruses has yet been formally identified in aphid vectors. A specialized region at the distal tip of aphid maxillary stylets, termed the acrostyle, exposes cuticular proteins, five of the six identified containing the conserved Rebers and Riddiford (R&R) consensus motif and belonging to the CPR_RR1 subfamily [20–23]. These proteins, Stylin-01, Stylin-02, Stylin-03, as well as the nearly identical Stylin-04 and Stylin-04bis (hereafter referred to as Stylin-04/04bis, 96% sequence identity), have been proposed as candidate receptors for stylet-borne viruses [22,23]. Two studies have been instrumental in identifying and validating the first receptor for cauliflower mosaic virus (CaMV, *Caulimovirus tessolobrassicae*) among these candidates. The RNAi-based study by Webster *et al.* [22], in which RNAi-mediated silencing of *Stylin-01* in aphids caused a marked reduction in CaMV transmission by the aphid species *Myzus persicae*, whereas silencing *Stylin-02* had no significant effect, demonstrating a specific role for Stylin-01 in CaMV transmission [22]. In the more recent work by Fu *et al.* [24], the use of CRISPR-Cas9 generated Stylin-01 mutants in *Acyrthosiphon pisum* genetically validated the role of Stylin-01 as the main receptor for CaMV [24]. A putative role of Stylin-01 or other stylins in potyvirus transmission has not yet been investigated, and the presence of potyvirus receptors at the acrostyle remains to be proven.

Here, we address these questions focusing on the turnip mosaic virus (TuMV, *Potyvirus rapae*), a potyvirus with the broadest known host range, causing significant damage to cruciferous crops in temperate and subtropical regions [25–27]. TuMV has been extensively studied over the last decades, and beyond its economic importance it remains a major model for studying virus-vector-plant interactions and more generally potyvirus biology [26]. We first characterized the TuMV retention sites in the stylets of three vector aphid species (*Aphis gossypii*, *M. persicae* and *A. pisum*) using complementary microscopy approaches. Recently generated and characterized stable *A. pisum* Stylin-01 mutant lines [24,28], in combination with RNAi-mediated silencing of *Stylin-04/04bis*, were used to characterize, for the first time, the role of stylins in TuMV transmission, providing new insights into the non-circulative transmission mechanism of potyviruses.

## Results

### Transmission of TuMV by three different aphid species

Numerous aphid species have been identified as vectors of TuMV [29]. To characterize TuMV receptors in aphid mouthparts, we first compared the ability of three aphid species – *A. gossypii*, *M. persicae* and *A. pisum* (wild-type, WT) – to transmit TuMV from infected turnip plants. Our results showed that the transmission efficiencies varied, with *A. gossypii* exhibiting the highest efficiency with a mean of 71.9 ± 2.0%, followed by *M. persicae* and *A. pisum* (WT), with means of 51.0 ± 4.3% and 39.6 ± 4.0% of transmission efficiencies, respectively, which are both significantly lower than *A. gossypii* (Fig 1).

**Fig 1.**
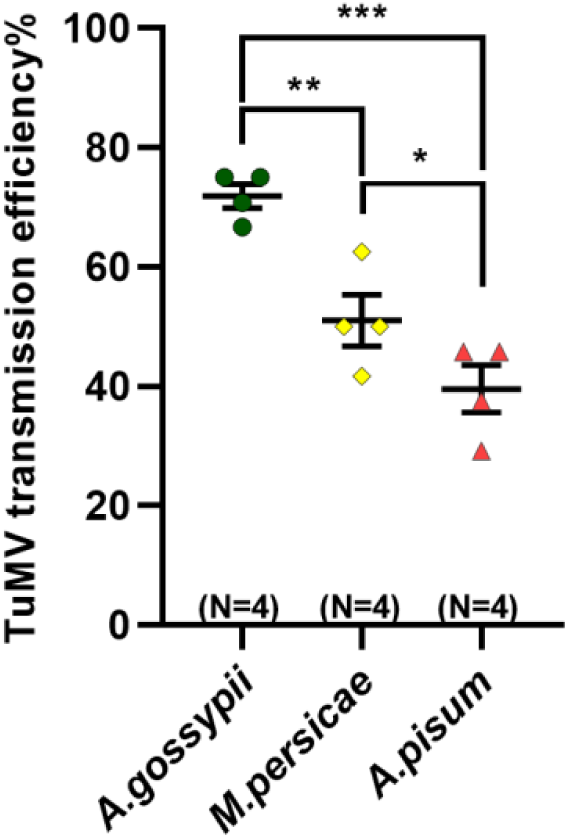
TuMV transmission efficiencies of three aphid species. The values obtained using one nymph per test plant are presented as mean ± SE of 4 independent replicates. Asterisks indicate significant differences according to generalized linear mixed models (GLMMs, \**P* < 0.05; \*\**P* < 0.01; \*\*\**P* < 0.001).

### TuMV retention sites in aphid stylets

Although the probable localization of retention sites in aphid stylets has been documented for a few potyviruses [6,7,30], the distribution of TuMV in the stylets of their vector has not yet been confirmed. Here, we adapted the protocol described by Mondal *et al.* [7] using an immunolabeling approach to detect TuMV particles in the stylets of the three aphid species fed on TuMV-infected turnips. Green fluorescence was observed exclusively in the stylets of aphids fed on infected turnips, with no signal detected in those fed on healthy plants (Fig 2A). Labeling appears in maxillary stylets as discontinuous punctate signals with variable intensity ranging from weak to strong. The number of labeled stylets among the three aphid species correlated with their TuMV transmission efficiencies, with *A. gossypii* showing the highest number of positive stylets (58.7 ± 3.4%), followed by *M. persicae* (50.7 ± 5.7%), while significantly fewer positive stylets were detected in *A. pisum* (WT) (41.9 ± 3.7%, *P* = 0.047) compared to *A. gossypii* (Fig 2B). Three labeling patterns were repeatedly observed across all three vector species: on the acrostyle, in the food canal at the distal tip of the maxillary stylets or in both the acrostyle and the food canal (Fig 2A). In the food canal, TuMV was not detected in a specific area but rather distributed in a zone above the acrostyle, and extending between 5 and 25 μm from the distal end of the stylets (Fig 2A). Notably, our results indicate that TuMV was detected mainly in the acrostyle region (either sole location or together with the food canal): the proportion of stylets labeled in the acrostyle being approximately 85% in *A. gossypii* (85.7 ± 7.9%) and in *M. persicae* (85.4 ± 7.3%), and slightly lower but not significantly different in *A. pisum* (WT, 68.3 ± 4.6%) (Fig 2C). In addition, the number of positive stylets in which TuMV exclusively binds to the acrostyle was significantly higher in *M. persicae* (57.9 ± 4.8%), compared to *A. pisum* (21.9 ± 9.1%, *P* = 0.042) and different but not significantly compared to *A. gossypii* (26.8 ± 7.7%, *P* = 0.075).

**Fig 2.**
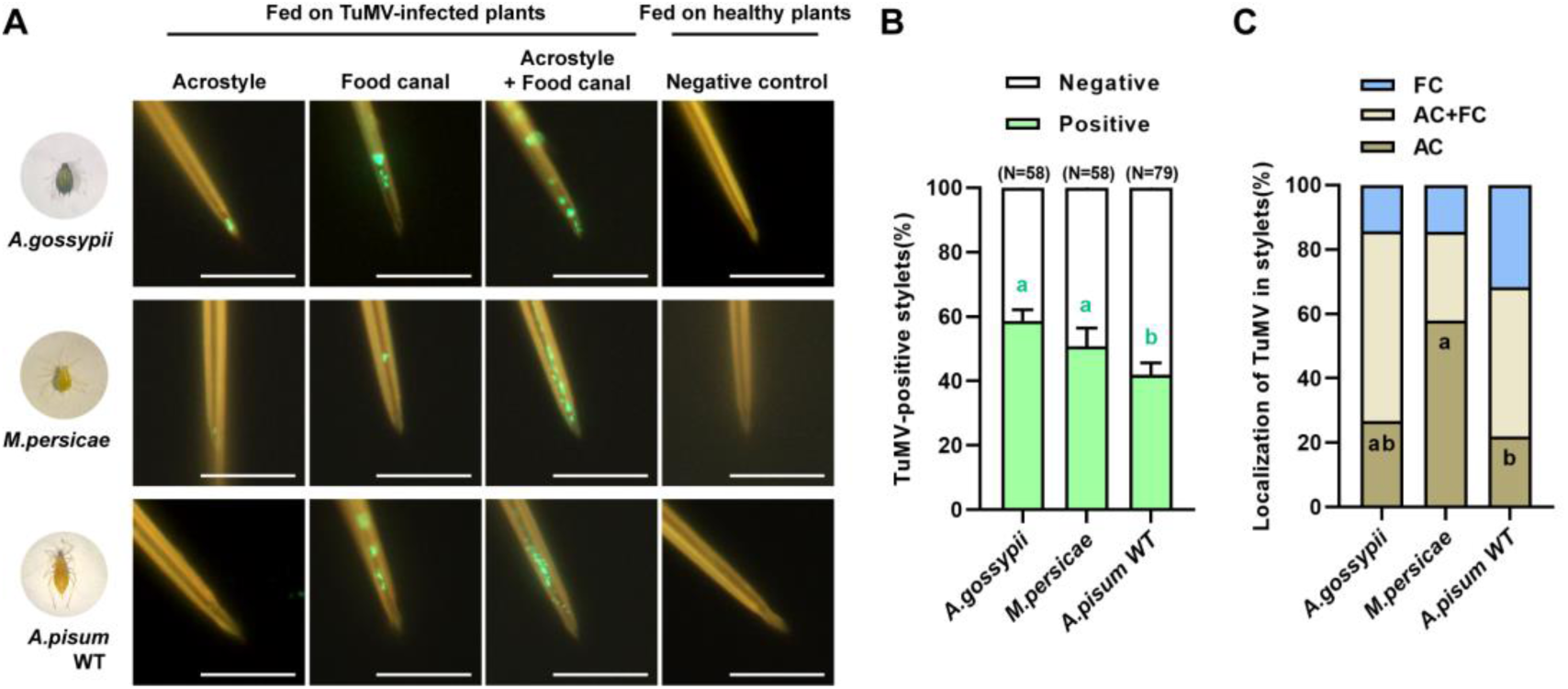
Detection of TuMV particles in the maxillary stylets of three aphid vector species by immunofluorescence labeling. (A) Immunofluorescence detection of TuMV particles in maxillary stylet of *A. gossypii, M. persicae* and *A. pisum* (WT) fed on infected turnips. Viruses appear as green fluorescence; aphids fed on healthy plants served as negative controls. (B) Proportion of TuMV-positive stylets among total maxillary stylets observed. “N” indicates the total number analyzed across three to five biological replicates. (C) Proportion of labeled stylets in the acrostyle (AC), food canal (FC), or both (AC + FC). (B and C) Different letters indicate significant differences according to generalized linear mixed models (GLMMs, *P* < 0.05). Scale bars in (A): 10 µm.

Alongside these observations, we also investigated TuMV retention sites in the stylets of *A. gossypii*, the best TuMV vector in our laboratory conditions, using transmission electron microscopy (TEM) according to the protocol developed by Uzest *et al.* [31] (Fig 3). We visualized virus particles in close association with the surface of the cuticle in the distal part of the food canal of maxillary stylets of *A. gossypii* fed on TuMV-infected turnips, near but not in the acrostyle region (Fig 3A-3C). These virus particles were visible as circular particles approximately of 12 nm in diameter in transverse orientation (Fig 3B and 3C), a shape and size consistent with the 3D model of the TuMV HC-Pro-virion transmissible complex (Fig 3D-3G; S1 and S2 Figs).

**Fig 3.**
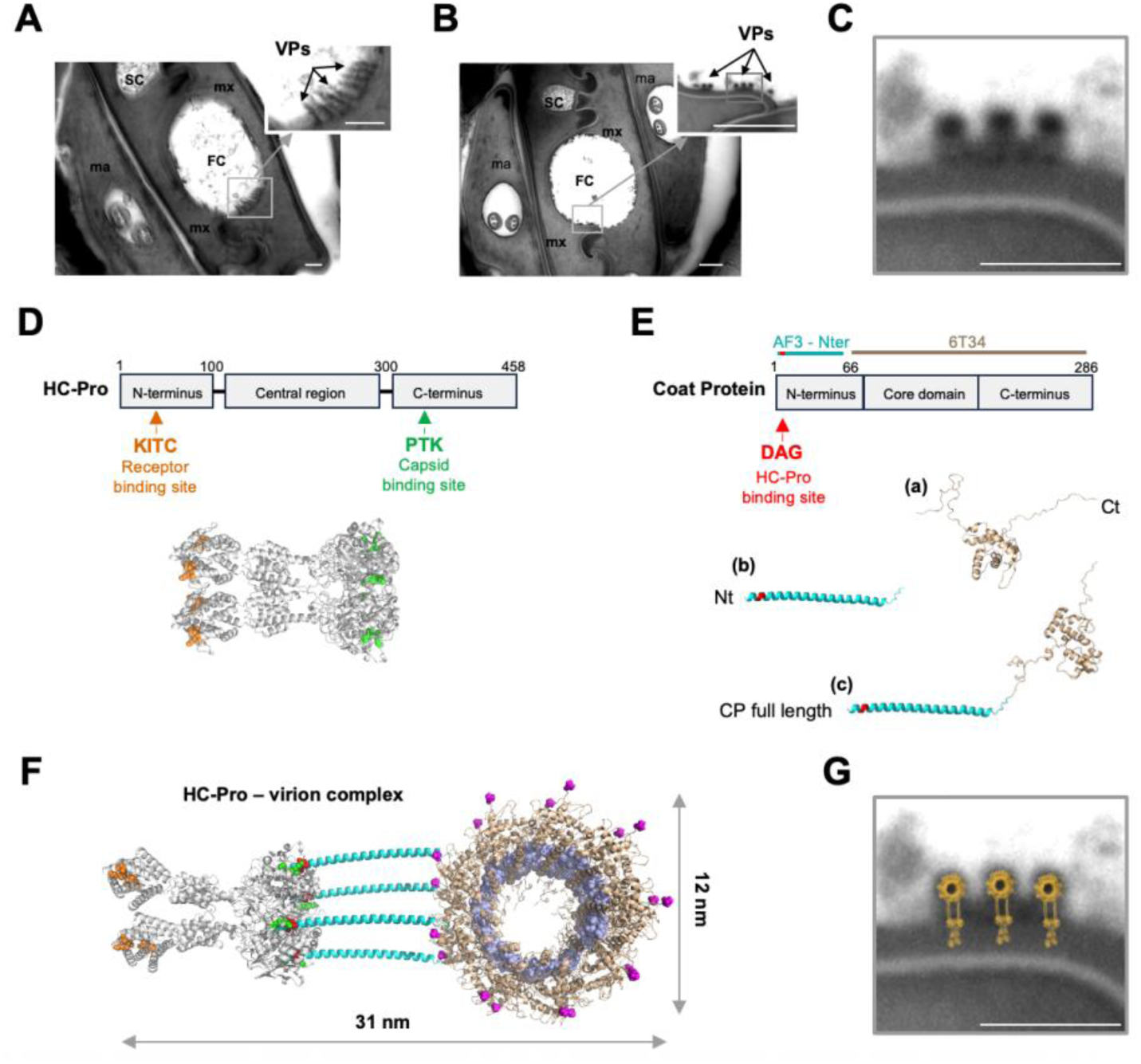
Observation of TuMV-like particles in the maxillary stylets of *A. gossypii* by transmission electron microscopy. (A-C) Representative electron micrographs of stylets cross-sections from *A. gossypii* after feeding on TuMV-infected turnip plants. Differences in the apparent morphology of the virus particles (VPs) between panels A and B likely reflect differences in the orientation of ultrathin cross-sections. This difference in sectioning angle is also reflected by the oblique appearance of the dendrites in the mandibular stylets. Insets show virus particles in the food canal of maxillary stylets (mx) in an apparent oblique (A) or near-transversal orientation (B). Mandibular stylets (ma) and salivary canal (SC) are indicated. (C) Enlargement of (B) showing three virus particles in close contact with the stylet cuticle. (D) AlphaFold3® (AF3) 3D model of TuMV HC-Pro tetramer based on previous 2D crystallography using electron microscopy studies (see Fig. S1 for AF3 prediction confidence). Domain organization is shown above; KITC (orange) and PTK (green) motifs involved in receptor and coat protein binding are indicated. (E) AF3-predicted TuMV coat protein (CP): (a) experimental partial solved structure lacking the 65 N-terminal amino acids (PDB ID:6T34, aa 66-270); (b) AF3 model of the N-terminal domain (aa 1-101) showing a large helix; (c) full-length AF3 model (aa 1-286); DAG (red) motif involved in HC-Pro binding is indicated (see Fig. S2 for prediction confidence). Domain organization is shown above. (F) AF3 model of the HC-Pro-virion transmissible complex showing one HC-Pro tetramer interacting with four CP N-terminal domains fitted into the TuMV virion structure in which the ssRNA molecule is indicated in violet (transverse view). (G) Three AF3 HC-Pro-virion complexes superposed onto the micrograph in C. Scale bars: 200 nm in (A) and (B), 50 nm in (C) and (G).

### Disruption of Stylin-01 impairs the capacity of aphids to transmit TuMV

The above results, indicating that TuMV binds mostly to the aphid acrostyle surface, suggest a potential role of stylin proteins in the binding process [23]. To determine whether Stylin-01, the receptor for CaMV, is also involved in TuMV transmission, we compared TuMV transmission capacity in *A. pisum*, the only aphid species with available mutants, across three cohorts: one wild-type and two Stylin-01 mutant lines. One of the mutant lines is a complete knockout, Sty01-KO, while the other produces a stylin mutated in its C-terminal part, Sty01-Cter [24,28]. TuMV transmission efficiency was markedly reduced in both mutant lines compared to the WT (Fig 4). The Sty01-KO mutant had the lowest transmission efficiency (23.6 ± 1.7%), representing a 53% decrease relative to WT (50.4 ± 2.3%, *P* < 0.0001). TuMV transmission was also reduced in Sty01-Cter aphids compared to WT (27.1 ± 1.8% transmission, *P* < 0.0001), but remained higher than in Sty01-KO aphids. Our results indicate that mutations in the C-terminal domain of Stylin-01 or its complete knockout impair aphids’ capacity to transmit TuMV. However, TuMV transmission remained substantial in both mutants, being only reduced by half. It has to be noted that the reduced TuMV transmission observed in the mutants was not attributable to defects in aphid feeding behavior. Electropenetrography analysis indeed revealed that WT and mutant aphids exhibited comparable feeding behavior patterns during the brief probing events associated with virus acquisition and inoculation (S3 Fig and S1 Table).

**Fig 4.**
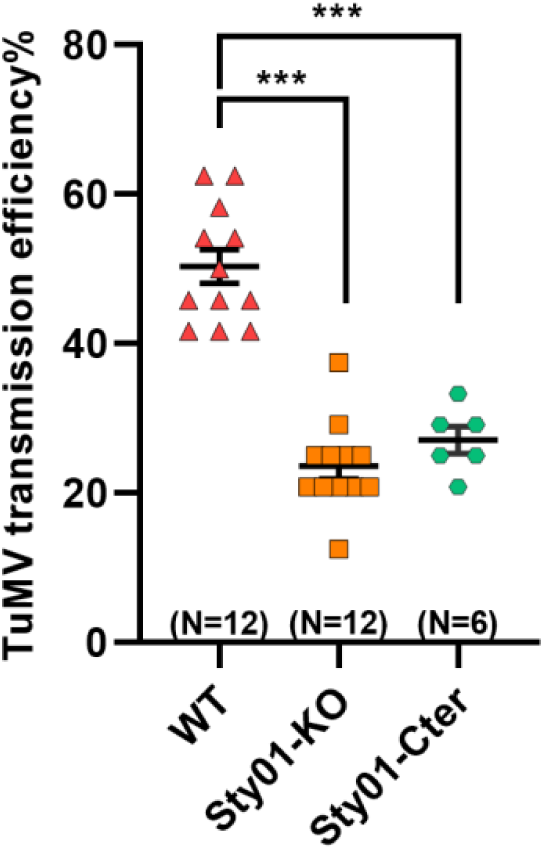
Impact of Stylin-01 mutations on TuMV transmission efficiency. The values obtained using two nymphs per test plant are presented as mean ± SE (N = 6-12 independent biological replicates). Asterisks indicate significant differences according to generalized linear mixed models (GLMMs, \*\*\**P* < 0.001).

### Stylin-01 mutations do not affect the distribution of TuMV in the stylets and acrostyle

To assess whether the presence of a mutated Stylin-01 or its absence affect the ability of the acrostyle to bind TuMV, we performed immunodetection of virus particles in the stylets of Sty01-KO and Sty01-Cter aphids fed on TuMV-infected plants (Fig 5A). Strikingly, the proportion of stylets detected as positive in the two mutant lines (50.9 ± 8.0% and 42.6 ± 3.6% in Sty01-KO and Sty01-Cter aphids, respectively) was not significantly different to that in the WT (41.9 ± 3.7%) (Fig 5B). Moreover, we observed that the distribution pattern of the virus in both mutant lines remained unchanged compared to the WT (Fig 5A), with TuMV detected in the acrostyle and food canal of mutant stylets. In addition, the proportion of stylets labeled at the acrostyle or both the acrostyle and food canal did not differ significantly between WT and mutant aphids (62.7 ± 17.3% of Sty01-KO stylets and 45.8 ± 7.1% of Sty01_Cter stylets) (Fig 5C).

**Fig 5.**
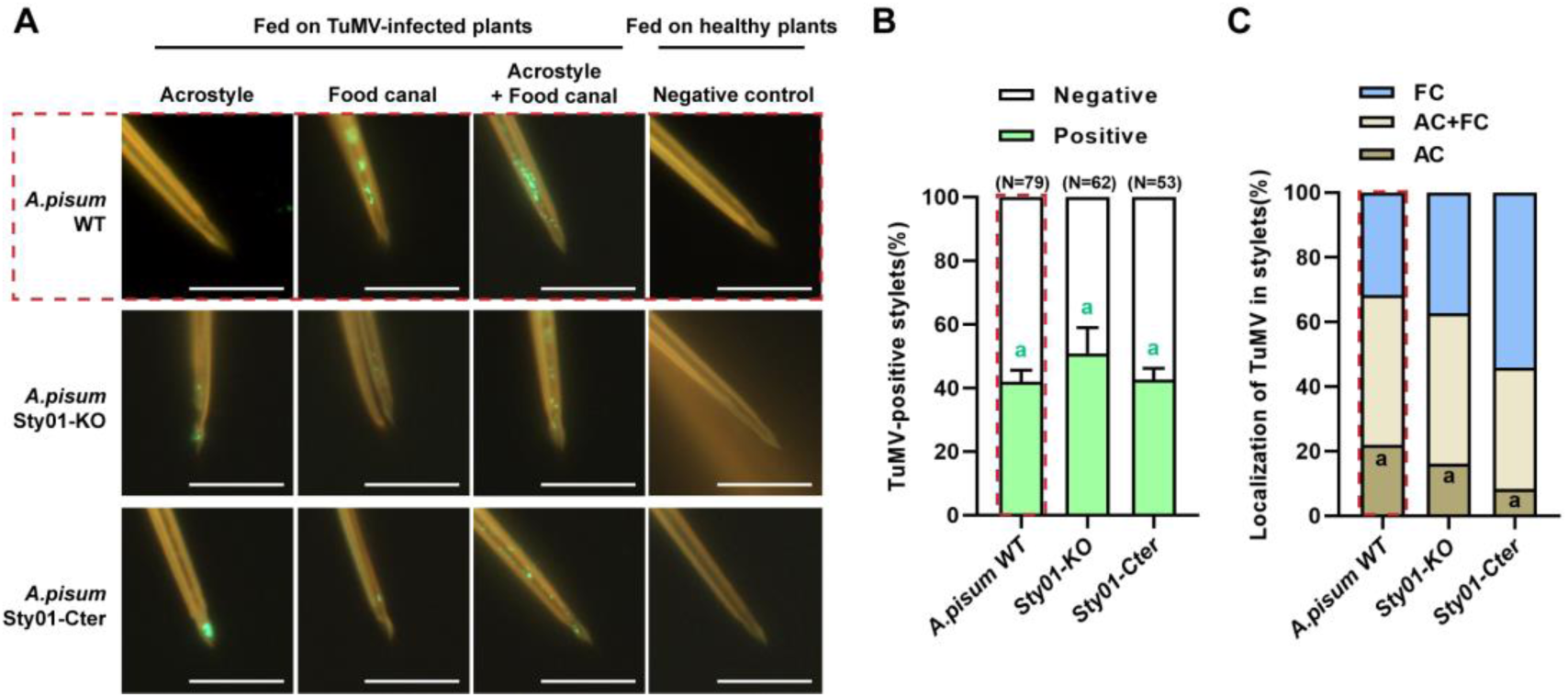
Detection of TuMV particles in the maxillary stylets of *A. pisum* WT and mutant lines by immunofluorescence labeling. (A) Detection of TuMV in the stylets of *A. pisum* WT, Sty01-KO and Sty01-Cter mutant lines fed on infected turnips using immunofluorescence labeling; aphids fed on healthy plants served as negative controls. (B) Proportion of TuMV-positive stylets in *A. pisum* WT and mutant lines. “N” indicates the total number analyzed across three to four biological replicates. (C) Proportion of labeled stylets in the acrostyle (AC), food canal (FC), or both (AC + FC) in *A. pisum* WT and mutant lines. All *A. pisum* WT data shown in this figure are identical to those presented Figure 2. They were incorporated to facilitate comparison with mutant lines and are delineated by red dashed lines for clarity. No significant differences according to generalized linear mixed models (GLMMs, *P* > 0.05). Scale bars in (A): 10 µm.

### Silencing *Stylin-04/04bis* reduces the capacity of aphids to transmit TuMV

Since Stylin-01 mutants still transmit TuMV with an efficiency of approximately 25%, and the virus is detected at comparable levels showing similar stylet distribution in both mutants and WT aphids, we hypothesized that, if Stylin-01 serves as a receptor for this virus, it is unlikely to be the sole receptor. To determine whether additional RR-1 stylins detected at the surface of aphid stylets are involved in TuMV transmission, we attempted to silence the *Stylin-02*, *03* and *04/04bis* genes in both WT and Sty01-KO backgrounds using specific siRNAs (sequences in S2 Table). However, we did not succeed in silencing the *Stylin-02* and *Stylin-03* genes in either aphid line under our experimental conditions (S4 Fig). Effective silencing was achieved only for the *Stylin-04/04bis* genes. In cohorts of first-instar nymphs fed on artificial diets supplemented with a specific siRNA targeting *Stylin-04/04bis* (siRNA-Sty04/04bis) for 24 or 48 h, the expression of *Stylin-04/04bis* was significantly reduced in WT nymphs when compared to that of nymphs from the control group fed on negative control siRNA (siRNA-NC) (*P_24h_* = 0.001, *P_48h_* = 0.047) (Fig 6A). We observed an optimal reduction of 39.1% of *Stylin-04/04bis* expression after 48 hours of treatment in WT nymphs. Similarly, *Stylin-04/04bis* transcripts were significantly reduced in Sty01-KO nymphs fed on artificial diets supplemented with specific siRNA-Sty04/04bis for 24 or 48 h when compared to control nymphs, with the highest reduction (50.9%) observed after 48 h of treatment (*P_24h_* = 0.006, *P_48h_* = 0.022) (Fig 6D). We verified that siRNA treatments did not affect aphid survival (S5 Fig). We therefore focused on the aphid phenotypes associated with *Stylin-04/04bis* gene silencing after a 48-hour exposure to artificial diets containing siRNAs. To achieve this, we subjected cohorts of WT and Sty01-KO aphids to siRNA ingestion, and analyzed one subset by qRT-PCR to evaluate silencing efficiency (Fig 6B and 6E), while using another one for TuMV transmission assays (Fig 6C and F). In WT aphids, when *Stylin-04/04bis* transcripts were reduced by 32.7% (Fig 6B), the TuMV transmission efficiency was significantly reduced by 53%, dropping from 58.9 ± 7.1% in controlled aphids to 27.8 ± 7.2% in silenced aphids (*P* < 0.001) (Fig 6C). The impact was even more substantial for Sty01-KO aphids for which a reduction of 36.9% of *Stylin-04/04bis* gene expression (Fig 6E) resulted in a 62% decrease in TuMV transmission efficiency between controlled and silenced aphids (42.1 ± 3.8% to 16.0 ± 2.5% of TuMV transmission efficiency, respectively; *P* = 0.002) (Fig 6F). We verified that the observed phenotype was not attributable to altered feeding behavior of silenced aphids during the transmission process (S6 Fig and S3 Table). No difference could be observed in the EPG variables related to TuMV acquisition or inoculation between silenced and control aphids.

**Fig 6.**
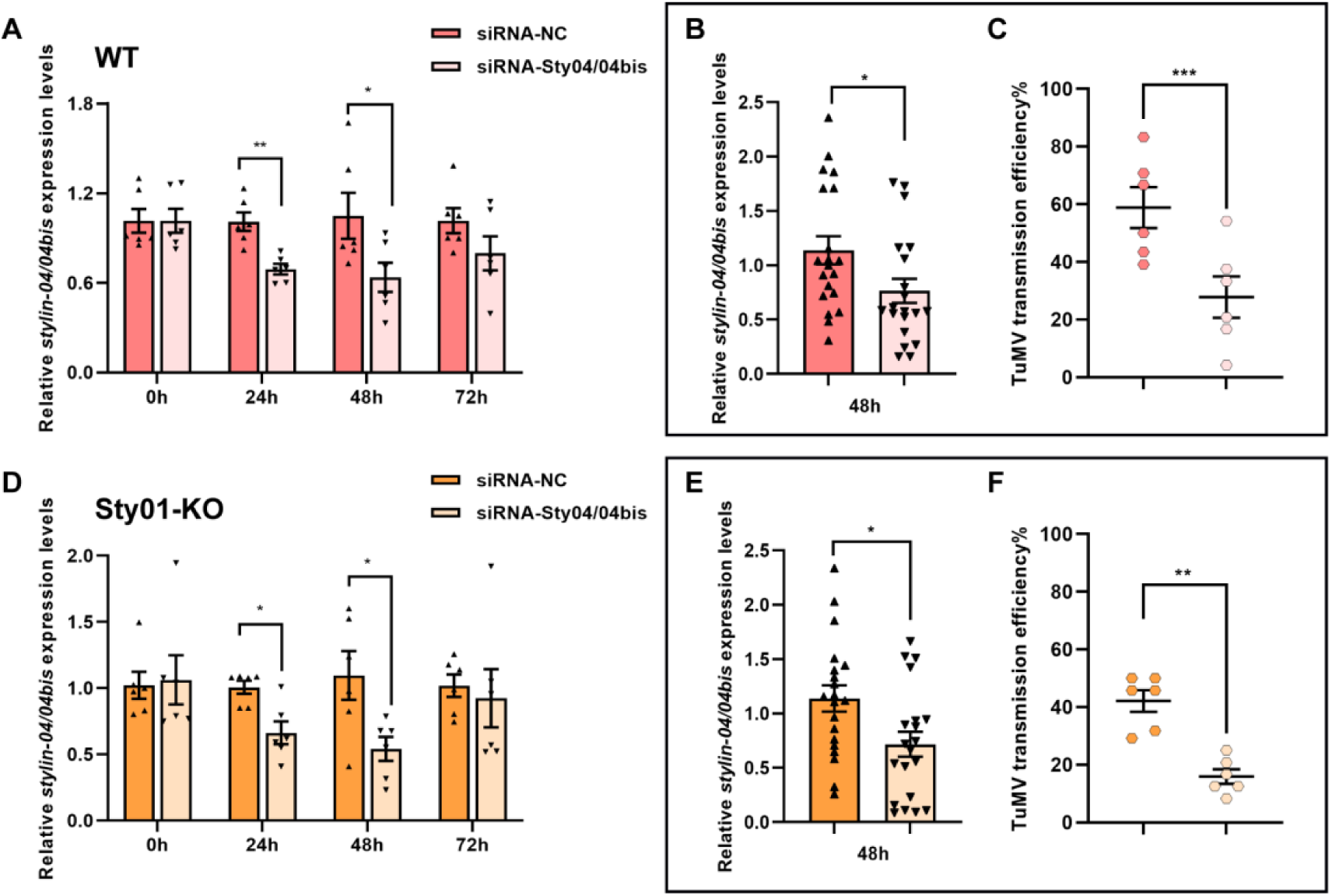
Effect of *Stylin-04/04bis* gene silencing on the TuMV transmission capacity of both *A. pisum* WT and Sty01-KO mutants. Relative expression levels of *Stylin-04/04bis* genes following RNAi treatment. First-instar WT (A-C) and Sty01-KO mutant (D-F) aphids were fed on artificial diets supplemented with gene-specific siRNAs targeting *Stylin-04/04bis* (siRNA-Sty04/04bis) or a negative control siRNA (siRNA-NC). Relative expression levels were determined by qRT-PCR using mitochondrial malate dehydrogenase (*Mdh2*) as the reference gene. (A, D) Expression levels at different time points after feeding (mean ± SE; N= 6 pools of 3 aphids). Cohorts of WT and Sty01-KO aphids were then subjected to 48 h of *Stylin-04/04bis* gene silencing treament and divided as follows: (B, E) A subset was analyzed by qRT-PCR to assess silencing efficiency (mean ± SE; N= 20 pools of 3 aphids). (C, F) The remaining individuals were used for TuMV transmission assays. Histograms show the proportion of TuMV-infected turnips after 3 h of exposure to two treated nymphs per plant (mean ± SE; 4 independent replicates). Statistical analyses were performed using Student’s *t*-test for gene expression data (\**P* < 0.05; \*\**P* < 0.01) and generalized linear mixed models (GLMMs) for transmission assays (\**P* < 0.05; \*\**P* < 0.01; \*\*\**P* < 0.001).

## Discussion

We previously demonstrated the essential role of cuticular stylin proteins in aphid stylets in the non-circulative transmission of caulimoviruses using CaMV as a model [22,24]. Here, we present novel results on the role of stylins in the transmission of the TuMV potyvirus, thereby extending our understanding of how stylet-borne viruses use these aphid cuticular proteins as virus receptors.

*A. pisum* is not generally considered a model vector for TuMV, unlike *M. persicae* and *A. gossypii* [32]; however, in order to use *A. pisum* mutants to evaluate the role of Stylin-01 in TuMV transmission, we first quantified its transmission efficiency to establish its suitability in our experimental systems. Our data revealed that *A. pisum* was the least effective vector for TuMV compared with *M. persicae* and *A. gossypii* (Fig 1). However, it showed a sufficiently high transmission rate to allow characterization of TuMV receptors in combination with the other two species.

We have thus proceeded to characterize retention sites for these three species in parallel. *In vitro* assays using either green fluorescent protein (GFP)-fused P2 or immunolabeling incubated directly on dissected aphid stylets, have enabled precise *in situ* mapping of CaMV receptors, demonstrating their exclusive distribution on the acrostyle surface [31]. By contrast, equivalent direct *in vitro* detection of the potyviral helper protein HC-Pro or potyviruses on dissected stylets has not been reported, and our own attempts to extend this approach to potyviruses have so far been unsuccessful. Potyvirus distribution in stylets has primarily been examined by transmission electron microscopy and, more recently, by immunofluorescence labeling following virus acquisition from infected plants or artificial diets [5–7,30]. However, these approaches provide limited spatial resolution and do not discriminate between virions retained at specific sites and those passively present in ingested contaminated fluid. Consequently, we need to keep in mind that fluorescence signals in stylets may not accurately reflect the precise localization of functional virus receptors. Despite these limitations, we have used a previously established immunofluorescence labeling [7] to visualize TuMV particles in the stylets of our three vector species. Similar to PVY [7], we detected TuMV particles on the distal part of the maxillary stylet. They were found at two different locations, the acrostyle and the food canal, in the 5-25 μm of the distal stylet tip (Fig 2A), in contrast with the single binding site of CaMV at the acrostyle. Although this approach does not allow us to assign functional relevance to each site unambiguously, the proportion of positive stylets across the three aphid species correlates with TuMV transmission efficiency, regardless of the distribution of detections across locations. This correlation suggests that all identified areas may contribute to virus retention (Fig 2C). However, since potyviruses are released during intracellular salivation [33,34], TuMV particles retained at the acrostyle or at the distal part of the food canal just above it are most likely to be effectively released and contribute to the subsequent infection.

We previously identified Stylin-01 as a key receptor mediating CaMV transmission. Its essential role has been demonstrated by the markedly reduced transmission efficiency observed in *M. persicae* cohorts following RNAi-mediated silencing of *stylin-01* [22], and, more recently, in an *A. pisum* CRISPR-Cas9 Stylin-01 knockout line, in which CaMV transmission capacity was nearly abolished [24]. We used the same CRISPR-edited aphid lines in the present study to examine whether Stylin-01 also contributes to TuMV transmission. Our data showed that both mutant lines exhibit significantly lower TuMV transmission capacity than the wild type. However, in contrast to what we observed for CaMV [24], TuMV transmission efficiency remained comparatively high, even in the KO mutant (Fig 4). Noticeably, although disruption of Stylin-01 reduces transmission efficiency of TuMV, it does not detectably affect its localization within the stylets. One possible explanation is that Stylin-01may also contribute to the structural organization of the acrostyle. In its absence, a redistribution of other stylins at the tip of maxillary stylets occurs, as observed recently [24], potentially modifying the spatial arrangement and accessibility of TuMV binding sites. Such changes may not be detectable through our standard immunolabeling approach but could impair the efficiency of acquisition, retention or inoculation. The reduced transmission efficiency observed for TuMV may reflect altered functional properties of the acrostyle rather than a complete loss of virus binding. Given that, in the mutants compared to the WT, the feeding behavior was unaffected during the phases related to TuMV acquisition and inoculation as assessed by EPG, and virus retention in aphid stylets remained unchanged, our combined results indicate that Sylin-01 contributes to TuMV transmission but is not strictly necessary for this process. The persistence of substantial transmission in the absence of Stylin-01 might be due to functional redundancy within the vector, with alternative stylet-associated proteins compensating for its loss.

Stylin-02, a paralog of Stylin-01, is overexpressed in Sty01-KO mutant aphids [24]. Stylin-02 is thought to partially compensate for the lack of Stylin-01 in CaMV retention as we detected this stylin over the entire acrostyle surface in Sty01-KO mutant, resulting in its co-localization with CaMV binding sites [24]. Based on these results, we hypothesize that Stylin-02 could play a role also in TuMV transmission in mutants, even though the correspondence between TuMV binding sites and Stylin-02, as for CaMV [24], cannot be clearly demonstrated. Unfortunately, we could not test this hypothesis in the present study due to the lack of RNAi inactivation efficacy in our experiments; therefore, this question warrants further investigation.

By contrast, with RNAi, we could test the potential role of *Stylin-04/04bis* by reducing gene expression by 40-50% in both WT and Sty01-KO aphids. For the RNAi experiments aphids had to be kept on control artificial diets for several days prior to transmission assays, and both lines exhibited slightly higher transmission efficiencies than in experiments conducted with aphids after they had fed on plants (Figs 4, 6C, and 6F). The fact that pre-feeding conditions (artificial diet versus plants) can influence aphid transmission efficiency has already been reported, particularly in the pea aphid, for which an increase in transmission of a circulative virus has been observed [35], but the underlying mechanisms for this phenomenon remain unresolved. Our study compares experiments conducted with wild-type and mutant strains under identical conditions using an artificial diet. In these experiments, even partial silencing of *Stylin04/04bis* resulted in a considerable reduction of TuMV transmission in both mutant aphid lines. We observed a more pronounced effect in the Sty01-KO line, with the silencing treatment leading to a 62% decrease in TuMV transmission, thereby highlighting a major role of both Stylin-01 and Stylin-04/04bis in this virus transmission process. Moreover, Stylin04/04bis was previously detected in the stylets of wild-type and Sty01-KO mutants at the apex of maxillary stylets with a weaker signal on the acrostyle edge and in its upper part in the vicinity of the food canal [23,24], a localization consistent with TuMV detection in pea aphid stylets (Fig 2A and 2D). The additive transmission defects observed upon disruption of Stylin-01 and Stylin-04/04bis reflects the contribution of these stylins to virus binding. The involvement of other stylins such as Stylin-02 and/or other cuticular proteins, in addition to Stylin-01 and Stylin-04/04bis, cannot be ruled out and would require further investigation, notably via CRISPR/Cas9 genome-editing approaches.

Consistent with the observed differences in the localization of the binding sites of TuMV and CaMV in *A. pisum* stylets, and with evidence that several stylins can assist the transmission of both viruses, a recent study showed no competition between TuMV and CaMV during acquisition by *Brevicoryne brassicae* [36]. When aphids acquired the viruses either simultaneously from mixed-infected plants or sequentially, transmission efficiencies remained unchanged regardless of acquisition order. These results support two non-exclusive scenarios. Either these viruses rely on distinct binding sites, with a major involvement of specific receptors in their transmission, implying a high level of specificity in virus-vector interactions; or they exploit alternative stylins thus enabling receptor substitution or compensation when the primary interaction is disrupted (as we observed in Stylin-01 KO mutants) or when binding sites are already occupied, thereby confering greater robustness and flexibility to virus-vector attachment. It would be interesting to determine whether Stylin-01 mutations also affect the transmission of other potyviruses. This question warrants further investigation as different viruses may rely on distinct interactions with aphid cuticular proteins.

In this work, we provide new insights into potyvirus receptors in aphid stylets. TuMV is detected in the acrostyle and in the distal part of the food canal, and our results reveal that aphid-mediated virus transmission relies on a multi-component interface including multiple stylins rather than a single receptor. Genetic disruption of Stylin-01 reduced transmission but did not abolish it, and partial silencing of Stylin-04/04bis impaired transmission in both wild-type and Stylin-01-deficient background, with greater effects in the knockout line. We demonstrate that TuMV requires several stylins to mediate its efficient transmission by aphids, with at least Stylin-01 and Stylin-04/04bis involved in the process. Receptor redundancy at the virus-vector interface may provide a more reliable binding platform for non-circulative plant viruses by allowing stylin overlapping to compensate for variability in the presence, abundance or accessibility of RR-1 cuticular proteins within and between vector species. Our study reveals the complex interaction between stylet-borne viruses and their insect vectors, providing a breakthrough in understanding the transmission mechanism of the largest group of plant RNA viruses.

## Materials and methods

### Aphid colonies, virus strain, and plants

Colonies of *Aphis gossypii* Glover and *Myzus persicae* Sulzer were maintained in a growth chamber at 23/18°C (day/night), with a photoperiod of 16/8 h (day/night) on *Cucurbita pepo* cv. Verte Noire Maraichère and *Solanum melongena* cv. Barbentane, respectively. Three *Acyrthosiphon pisum* lines were used in this study: one wild-type line (WT) and two Stylin-01 mutant lines: Sty01-KO (complete gene knockout) and Sty01-Cter (only Cter deletion) [28]. All three lines displayed similar developmental time and no detectable growth abnomalities, as previuously reported [24]. They were maintained in controlled conditions at 18°C, with a 16/8 h (day/night) photoperiod, on faba bean (*Vicia faba* cv. Sutton). TuMV (isolate C42) was maintained on turnips *Brassica rapa* cv. Just Right by mechanical inoculation. Healthy and infected turnips were maintained at 23/18°C under a 16/8 h (day/night) photoperiod.

### Aphid transmission tests

*Assays to compare TuMV transmission efficiencies of three different aphid species A. gossypii* and *M. persicae* adults and *A. pisum* (WT) N1 aphids (one-day-old nymphs post synchronization) were collected and starved for 1 h in a glass tube prior to transmission tests. They were then allowed a 2-min acquisition access period (AAP) on the 8^th^ leaf of a TuMV-infected turnip used as a virus source plant 18 days post-inoculation. Aphids were immediately collected in a glass petri dish and transferred to 8-day-old turnip test plants for a 3-h inoculation access period (IAP) followed by insecticide treatment. The following design was used in the experiments: 24 test plants per source plant and 1 aphid per test plant. Aphid transmission efficiency was calculated by recording, 21 days post-transmission, the number of test plants showing viral symptoms out of the total number of test plants used. Four independent biological replicates were performed for all aphid species.

#### Assays to compare TuMV transmission efficiencies across A. pisum lines

Two N2 aphids (2-day-old nymphs post synchronization) were used per test plant to compare the TuMV transmission efficiencies of *A. pisum* WT and mutant lines, Sty01-KO and Sty01-Cter, as well as to compare the transmission efficiencies of WT and Sty01-KO silenced nymphs. At least six independent biological replicates were performed for all aphid lines and all treatments.

### *In vitro* interaction assays on dissected stylets

Detection of TuMV on aphid stylets was performed according to a protocol described by Mondal *et al.* [7] with some modifications. *A. gossypii*, *M. persicae* and *A. pisum* adults were starved for 2 h prior to a 2-min acquisition access period (AAP) on the 8^th^ leaf of either healthy or TuMV-infected turnip at 21 days post-inoculation (dpi). Immediately after feeding, aphids were transferred into a tube immersed in liquid nitrogen and subsequently placed in fixation buffer (4% paraformaldehyde in phosphate-buffered saline buffer, PBS) until dissection. Stylet bundles were excised from aphid heads using fine tweezers and incubated in 0.1% Triton-X100 solution for 1 h at room temperature. Samples were washed three times with PBS supplemented with 0.5% Tween 20 (PBST buffer), followed by incubation for 1 h in blocking buffer (1% bovine serum albumin in PBST buffer). Stylet bundles were incubated overnight at 4°C with primary TuMV coating antibody (CAB 18700/1000, Agdia EMEA, Soisy-sur-Seine, France) at a 1:200 dilution. After washing, samples were incubated with Alexa Fluor 488-conjugated goat anti-rabbit IgG secondary antibody (A11070, Thermo Fisher Scientific, Waltham, MA) at a 1:400 dilution for 4 h at room temperature. Finally, stylets were washed in PBST, and maxillary and mandibular stylets were separated and then mounted in Mowiol mounting medium on glass slides. Images were acquired with a confocal laser scanning microscope (ZEISS LSM900). At least three independent biological replicates were performed with a minimum of 15 stylets observed for each experiment.

### Transmission electron microscopy

Detection of TuMV particles in aphid stylets by transmission electron microscopy was conducted on *A. gossypii* as it is the best TuMV vector in our laboratory conditions. Adult aphids fed for 2 min on a systemically TuMV-infected turnip were removed carefully and immediately anesthetized with CO_2_. Their heads were removed and fixed in a 4% glutaraldehyde - 0.1 M sodium cacodylate fixation buffer (FB) pH 7.2 for 4 h at 4°C according to Uzest et al. [31]. Samples were rinsed in FB buffer before post-fixation in 1% osmic acid in FB for 1 h at 4°C. They were then dehydrated in a graded series of acetone up to 100% and finally embedded in epoxy resin (TAAB 812). Sections 60-nm thick were observed in a JEOL JEM 1400 microscope.

### Analysis of aphid feeding behavior

The same experimental design used for the transmission test was applied to perform Electrical Penetration Graph (EPG) recordings to monitor the feeding behavior of N2 instar nymphs during acquisition and ingestion activities associated with TuMV transmission, following the protocol described previously [24]. Feeding behavior was recorded using a Giga-8 DC-EPG device (EPG Systems, Wageningen, The Netherlands) for 10 min from the start of probing. This duration was sufficient to capture all waveforms associated with the Acquisition Access Period (AAP) and Inoculation Access Period (IAP) required for the transmission of non-circulative, non-persistent viruses [37]. After the 10-min feeding period on a TuMV-infected plant, the wired aphids were immediately removed and transferred to a young, healthy turnip test plant, thereby replacing the infected source plant. EPG recording was then resumed. Each plant was used only once and replaced for each aphid. At least 22 aphid nymphs were recorded per aphid line. The EPG data were acquired and analyzed using Stylet+ software for Windows (EPG Systems). Signals were analyzed manually using the software A2-EPG [38] focusing on the following variables: the time from the beginning of the first probe to first intracellular puncture characterized by a potential drop (t > 1pd), the duration of the first intracellular puncture (d_1pd), the duration of subphases II-1, II-2 and II-3 during the first intracellular puncture (d_1pd II-1, d_1pd II-2 and d_1pd II-3, respectively), II-3 being associated with TuMV acquisition and and II-1 with TuMV inoculation [39–41].

### RNA interference

The negative control siRNA (siRNA-NC; SR-CL000-005, Eurogentec) and siRNAs specifically targeting *Stylin-02*, or *Stylin-03* or *Stylin-04/04bis* mRNA (sequences provided in Table S2) were mixed with AP3 sterile artificial diet [42] at a final concentration of 1 µM. The mixture was orally delivered to synchronized first-instar nymphs (N1) (< 4-h-old) through a Parafilm membrane sachet. Fifty nymphs per treatment were transferred to each sachet containing 400 µL of diet and maintained at 18°C for 24, 48, or 72 h. For gene expression analysis, 3 to 10 pools of 3 surviving aphids per treatment were collected. For TuMV transmission assays and EPG experiments, nymphs were collected after 48 h of artificial feeding.

### Reverse-transcription quantitative PCR

Total RNA was extracted using a RNeasy mini kit (Qiagen, Hilden Germany) and treated with RQ1 RNAse-free DNAse I (Promega Corporation, Madison, WI). First-strand cDNA was synthesized from 200 ng of total RNAs using Moloney murine leukemia virus (MMLV) reverse transcriptase (Promega Corporation, Madison, WI) and oligo (dT) as a primer according to the manufacturer’s instructions. All quantitative PCRs were performed in triplicates on a LightCycler 480 instrument using a LightCycler 480 SYBR green I master mix (Roche, Penzberg, Germany) according to the manufacturer’s recommendations with gene-specific primers (S4 Table). The gene encoding the mitochondrial malate dehydrogenase (Mdh2) was used as reference gene [24]. Relative expressions were determined by using the 2-ΔΔCT method [43].

### Structure prediction

The structural models of the TuMV HC-Pro protein, the TuMV CP protein, and HC-Pro-viron complexes of TuMV were created using Alphafold 3 ® [44].

### Statistical analysis

Statistical analyses were performed using SPSS Statistics 20.0 software, and plotted using GraphPad Prism version 9. Differences in aphid transmission efficiencies, stylets labeling and aphid survival were tested using generalized linear mixed models (GLMMs). Differences in stylin genes expression and aphid survival rate was analyzed using Student’s *t*-test. Differences in EPG variables were analyzed using Kruskal-Wallis or Mann-Whitney U tests. Asterisks denoted statistical significance between two groups (\**P* < 0.05, \*\**P* < 0.01, and \*\*\**P* < 0.001). Identical letters indicate differences among groups that are statistically nonsignificant.

## Supporting Information

**S1 Fig. AlphaFold3® 3D model of a tetramer of HC-Pro of the turnip mosaic virus (TuMV, *Potyvirus rapae*).** (A) C-terminal region of TuMV HC-Pro, which contains the PTK motif interacting with the viral coat protein, tetrameric state based on previous 2D crystallography using electron microscopy studies [14]. This model and his tetrameric state were predicted with high accuracy (interface predicited TM-score ipTM, and predicted TM-score pTM > 85). (B) TuMV HC-Pro full-length showing the tetrameric state of the C-terminal region and an N-terminal domain (aa 1-170) formed by two dimers. A good but not optimal prediction for the N-terminal part (aa 1–100, light blue) comprising the KITC motif involved in the interaction with the receptor in the insect vector; and a poor prediction for the central region (aa 100–170, yellow).

**S2 Fig. AlphaFold3® 3D model of the TuMV coat protein (TuMV CP) N-terminal domain (aa 1-101) and in full-length (aa 1–286).** (A) The N-terminal domain of TuMV CP (aa 1-101), which is missing from the PDB ID :6T34 3D structure, forms a large helix with good level of confidence. (B) AF3 model of TuMV CP full-length protein (aa 1-286). The regions are colored according to predicted Local Distance Difference Test pLDDT values.

**S3 Fig. Feeding behavior of wild-type and mutant A. pisum lines during TuMV acquisition and inoculation.** Short-term feeding behavior of wild-type and mutant nymphs was monitored by electrical penetration graph (EPG) analysis using the protocol previously described for assessing the role of stylin-01 in CaMV transmission [24]. EPG variables associated with virus acquisition (A) or inoculation (B) were analyzed (see Table S1). The first intracellular puncture (pd), identified by a characteristic potential drop in EPG recordings, is critical for potyvirus transmission. Virus acquisition occurs during aphid ingestion in pd subphase II-3, whereas inoculation is associated with salivation during pd subphase II-1 [33,34]. (A) Variables related to TuMV acquisition: time from the beginning of the first probe to the first intracellular puncture (t > 1pd) and duration of sub-phase II-3 within the first pd (d_1pd II-3) (N = 22–25, Kruskal-Wallis test; ns, not significant). (B) Variables related to TuMV inoculation: t > 1pd and duration of sub-phase II-1 within the first pd (d_1pd II-1) (N = 22–25, Kruskal-Wallis test; ns, not significant). In both panels, aphids plotted below the dotted line reached the first intracellular puncture within 30 s of recording onset. No significant difference was observed among the three cohorts in the time required to reach the first punctured cell on TuMV-infected plants (acquisition) or healthy plants (inoculation).

**Fig S4. Gene silencing efficiency of siRNAs targeting *Stylin-02* or *Stylin-03* expression in *A. pisum* WT and Sty01-KO mutants.** Relative expression levels of *Stylin-02* and *Stylin-03* genes at different time points following RNAi treatment (mean ± SE; N= 3–10 pools of 3 aphids). First-instar WT (A-B) and Sty01-KO (C-D) aphids were fed artificial diets supplemented with gene-specific siRNAs targeting *Stylin-02* (siRNA-Sty02), *Stylin-03* (siRNA-Sty03), or with a negative control siRNA (siRNA-NC). Relative expression levels were determined by qRT-PCR using mitochondrial malate dehydrogenase (*Mdh2*) as the reference gene. (A) *Stylin-02* gene expression levels in WT treated aphids. (B) *Stylin-03* gene expression levels in WT treated aphids. (C) *Stylin-02* gene expression levels in Sty01-KO treated aphids. (D) *Stylin-03* gene expression levels in Sty01-KO treated aphids. No significant difference in expression was observed according to Student’s *t*-test (*P* > 0.05).

**S5 Fig. Effect of *Stylin-04/04bis* gene silencing on the survival of *A. pisum* WT and Sty01-KO mutants.** Survival of aphids after 48 h of feeding on artificial diets supplemented with gene-specific siRNAs. Cohorts of WT (A) and Sty01-KO mutant (B) aphids were fed on artificial diets supplemented with siRNAs targeting *Stylin-04/04bis* (siRNA-Sty04/04bis) or a negative control siRNA (siRNA-NC). Bars show the minimum and maximum values, (+) indicate the mean of 20 independent replicates. Statistical analysis was assessed using generalized linear mixed models (GLMMs); ns, not significant (*P* > 0.05).

**S6 Fig. Feeding behavior of wild-type and Sty01-KO mutant *A. pisum* lines during TuMV acquisition and inoculation after silencing *stylin-04/04bis*.** Short-term feeding behavior of wild-type and mutant nymphs fed for 48 hours on artificial diets supplemented with specific siRNA targeting *Stylin04/04bis* expression (siRNA-Sty04/04bis) or negative control siRNA (siRNA-NC) was monitored by electrical penetration graph (EPG) analysis. (A-E) Data retrieved from WT treated aphids. (F-J) Data retrieved from Sty01-KO treated aphids. Relative *Stylin-04/04bis* expression levels of WT (A) and Sty01-KO aphids (F) exposed for 48 hours to siRNA artificial diets (mean ± SE; N = 6 pools of 3 aphids; asterisks show significant differences according to Student’s *t*-test, \**P* < 0.05; \*\**P* < 0.01). First-instar nymphs were used at the start of the experiment. EPG variables associated with acquisition on infected turnips (B-C, G-H) or inoculation on healthy plants (D-E, I-J) were analyzed (see Table S3). Variables related to TuMV acquisition: (B, G) time from the beginning of the first probe to the first intracellular puncture (t > 1pd) and (C, H) duration of sub-phase II-3 within the first pd (d_1pd II-3) (N = 26-27, Mann-Whitney U test; ns, not significant). Variables related to TuMV inoculation: (D, I) t > 1pd and (E, J) duration of sub-phase II-1 within the first pd (d_1pd II-1) (N = 26-27, Mann-Whitney U test; ns = not significant). In panels (B, D, G, I), aphids plotted below the dotted line reached the first intracellular puncture within 30 s of recording onset. No significant difference was observed in the feeding behaviors associated with TuMV acquisition on infected plants or TuMV inoculation on healthy plants, regardless of the treatment and aphid lines.

**S1 Table. Comparison of the feeding behavior of N2 nymphs of WT and mutant *A. pisum* lines.** Feeding behavior recorded for 10 min on turnip plants infected with TuMV (21 dpi) for EPG variables related to virus acquisition, and for 10 min on young healthy turnip plants for EPG variables related to virus inoculation. Time is expressed in seconds (mean ± SE).

**S2 Table. List of oligonucleotide siRNAs used in this study to silence stylin genes.**

**S3 Table. Comparison of the feeding behavior of N2 nymphs of WT and Sty01-KO lines after silencing *Stylin-04/04bis.*** Feeding behavior recorded for 10 min on turnip plants infected with TuMV (21 dpi) for EPG variables related to virus acquisition, and for 10 min on young healthy turnip plants for EPG variables related to virus inoculation. Time is expressed in seconds (mean ± SE).

**S4 Table**. **List of the oligonucleotide primers used for qRT-PCR analyses.**

## Acknowledgements

We are grateful to Juan Jose Lopez-Moya, Stéphane Blanc and Manuella van Munster for fruitful discussions. This work was supported by grants from the French National Research Agency (ANR-15-CE20-0011 and ANR-21-CE20-0001) and from the Bill and Melinda Gates Foundation (GCEag, OPP1130147). Yu Fu acknowledges the Chinese Scholarship Council (CSC Grant No. 202103250004) and the French National Institute Research for Agriculture, Food and Environment INRAE, for financial support of her PhD studies. All rearing and experiments on insects have been performed on the Baillarguet Vectoplante platform, member of the Vectopole Sud Network.

