## Supplementary material for "Aphid stylin cuticular proteins contribute to turnip mosaic virus (*Potyvirus*) transmission": Fu-et-al_Supporting-Information

**Fig. S1.** AlphaFold3® 3D model of a tetramer of HC-Pro of the turnip mosaic virus (TuMV, *Potyvirus rapae*).

**Fig. S2.** AlphaFold3® 3D model of the TuMV coat protein (TuMV CP) N-terminal domain (aa 1-101) and in full-length (aa 1–286).

**Fig. S3.** Feeding behavior of wild-type and mutant *A. pisum* lines related to TuMV acquisition and inoculation.

**Fig. S4.** Gene silencing efficiency of siRNAs targeting *Stylin-02* or *Stylin-03* expression in *A. pisum* WT and Sty01-KO mutants.

**Fig. S5.** Effect of *Stylin-04/04bis* gene silencing on the survival of *A. pisum* WT and Sty01-KO mutants.

**Fig. S6.** Feeding behavior of wild-type and Sty01-KO mutant *A. pisum* lines related to TuMV acquisition and inoculation after silencing *Stylin-04/04bis*.

**Table S1.** Comparison of the feeding behavior of N2 nymphs of WT and mutant *A. pisum* lines.

**Table S2.** List of oligonucleotides used in this study to silence *Stylin* genes.

**Table S3.** Comparison of the feeding behavior of N2 nymphs of WT and Sty01-KO lines after silencing *Stylin-04/04bis.*

**Table S4**. List of the oligonucleotides used for qRT-PCR analyses.

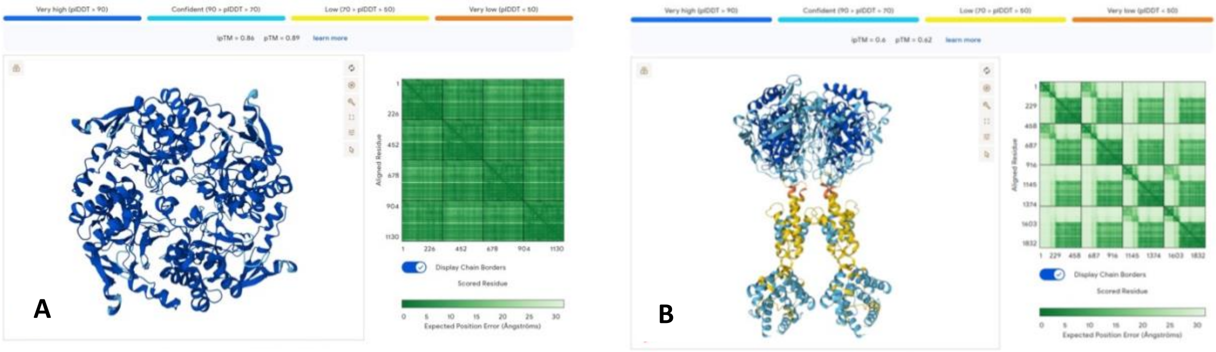


**Fig S1. AlphaFold3® 3D model of a tetramer of HC-Pro of the turnip mosaic virus (TuMV, *Potyvirus rapae*).**
(A) C-terminal region of TuMV HC-Pro, which contains the PTK motif interacting with the viral coat protein, tetrameric state based on previous 2D crystallography using electron microscopy studies (Plisson et al., 2003). This model and his tetrameric state were predicted with high accuracy (interface predicited TM-score ipTM, and predicted TM-score pTM > 85). (B) TuMV HC-Pro full-length showing the tetrameric state of the C-terminal region and an N-terminal domain (aa 1-170) formed by two dimers. A good but not optimal prediction for the N-terminal part (aa 1–100, light blue) comprising the KITC motif involved in the interaction with the receptor in the insect vector; and a poor prediction for the central region (aa 100–170, yellow).

Plisson C, Drucker M, Blanc S, German-Retana S, Le Gall O, Thomas D, Bron P. 2003. Structural characterization of HC-Pro, a plant virus multifunctional protein. J Biol Chem. 2003 Jun 27;278(26):23753-61. doi: 10.1074/jbc.M302512200.

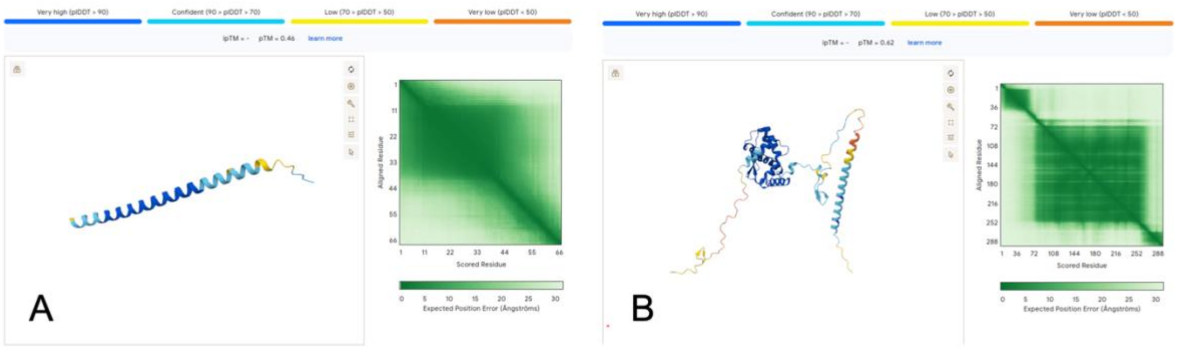


**Fig S2. AlphaFold3® 3D model of the TuMV coat protein (TuMV CP) N-terminal domain (aa 1-101) and in full-length (aa 1–286).**

(A) The N-terminal domain of TuMV CP (aa 1-101), which is missing from the PDB ID :6T34 3D structure, forms a large helix with good level of confidence. (B) AF3 model of TuMV CP full-length protein (aa 1-286). The regions are colored according to predicted Local Distance Difference Test pLDDT values.

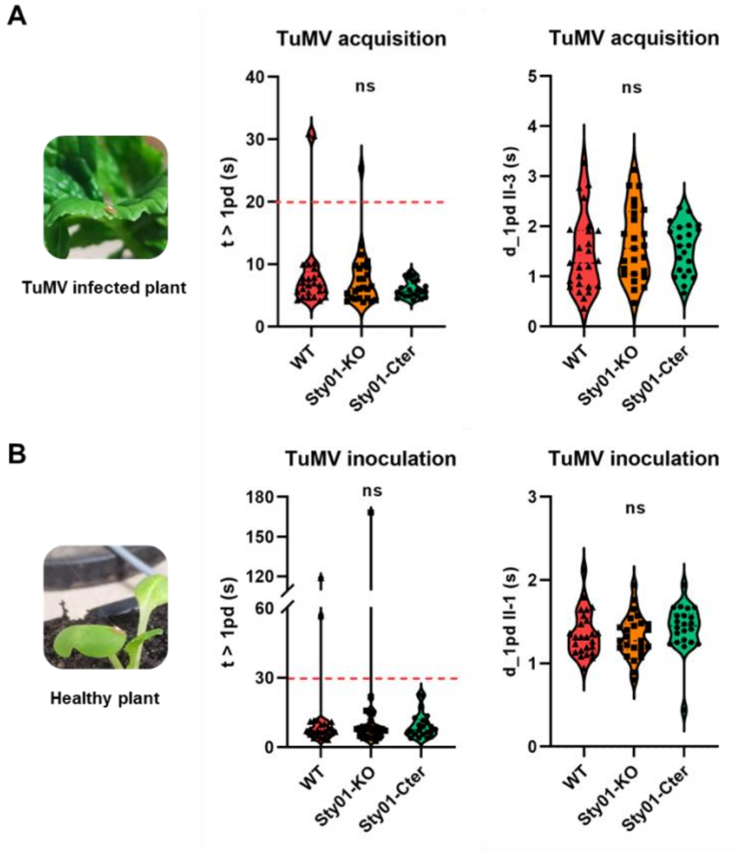


**Fig S3. Feeding behavior of wild-type and mutant *A. pisum* lines during TuMV acquisition and inoculation.**

Short-term feeding behavior of wild-type and mutant nymphs was monitored by electrical penetration graph (EPG) analysis using the protocol previously described for assessing the role of stylin-01 in CaMV transmission (Fu et al., 2025). EPG variables associated with virus acquisition (A) or inoculation (B) were analyzed (see Table S1). The first intracellular puncture (pd), identified by a characteristic potential drop in EPG recordings, is critical for potyvirus transmission. Virus acquisition occurs during aphid ingestion in pd subphase II-3, whereas inoculation is associated with salivation during pd subphase II-1 (Martin et al., 1997; Powell, 2005). (A) Variables related to TuMV acquisition: time from the beginning of the first probe to the first intracellular puncture (t > 1pd) and duration of sub-phase II-3 within the first pd (d_1pd II-3) (N = 22–25, Kruskal-Wallis test; ns, not significant). (B) Variables related to TuMV inoculation: t > 1pd and duration of sub-phase II-1 within the first pd (d_1pd II-1) (N = 22–25, Kruskal-Wallis test; ns, not significant). In both panels, aphids plotted below the dotted line reached the first intracellular puncture within 30 s of recording onset. No significant difference was observed among the three cohorts in the time required to reach the first punctured cell on TuMV-infected plants (acquisition) or healthy plants (inoculation).

Fu Y, Deshoux M, Cayrol B, Le Blaye S, Achard E, Hudaverdian S, Cloteau R, Pichon E, Strozyk E, Prunier-Leterme N, Jousselin E, Sauvion N, Thébaud G, Le Trionnaire G, Colella S, Uzest M. 2025. Stylet cuticular gene-directed mutagenesis impairs the pea aphid vector capacity to transmit a plant virus. PLoS Pathog. 2025 May 23;21(5):e1013192. doi: 10.1371/journal.ppat.1013192.

Martín, B., Collar, J.L., Tjallingii, W.F. and Fereres, A. 1997. Intracellular ingestion and salivation by aphids may cause the acquisition and inoculation of non-persistently transmitted plant viruses. Journal of General Virology, 78, 2701–2705.

Powell, G. 2005. Intracellular salivation is the aphid activity associated with inoculation of non-persistently transmitted viruses. The Journal of General Virology, 86, 469–472.

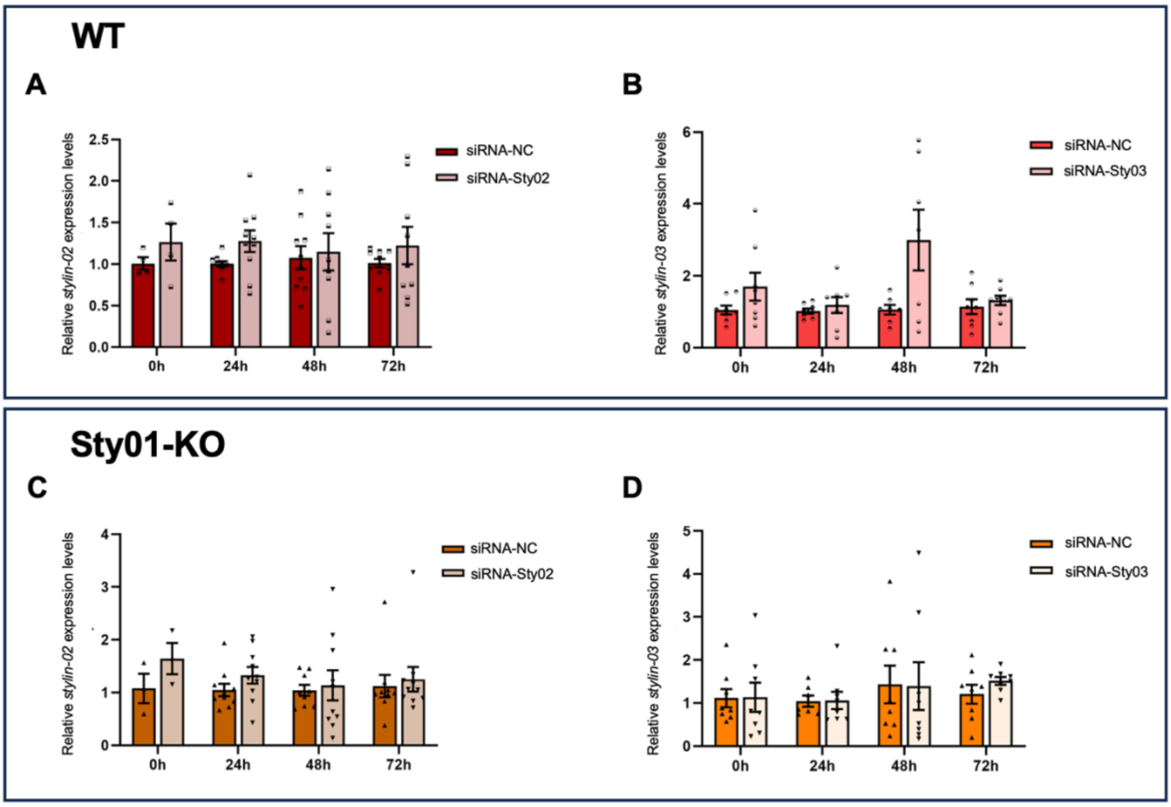


**Fig S4. Gene silencing efficiency of siRNAs targeting *Stylin-02* or *Stylin-03* expression in *A. pisum* WT and Sty01-KO mutants.**

Relative expression levels of *Stylin-02* and *Stylin-03* genes at different time points following RNAi treatment (mean ± SE; N= 3–10 pools of 3 aphids). First-instar WT (A-B) and Sty01-KO (C-D) aphids were fed artificial diets supplemented with gene-specific siRNAs targeting *Stylin-02* (siRNA-Sty02), *Stylin-03* (siRNA-Sty03), or with a negative control siRNA (siRNA-NC). Relative expression levels were determined by qRT-PCR using mitochondrial malate dehydrogenase (*Mdh2*) as the reference gene. (A) *Stylin-02* gene expression levels in WT treated aphids. (B) *Stylin-03* gene expression levels in WT treated aphids. (C) *Stylin-02* gene expression levels in Sty01-KO treated aphids. (D) *Stylin-03* gene expression levels in Sty01-KO treated aphids. No significant difference in expression was observed according to Student's *t*-test ($P > 0.05$).

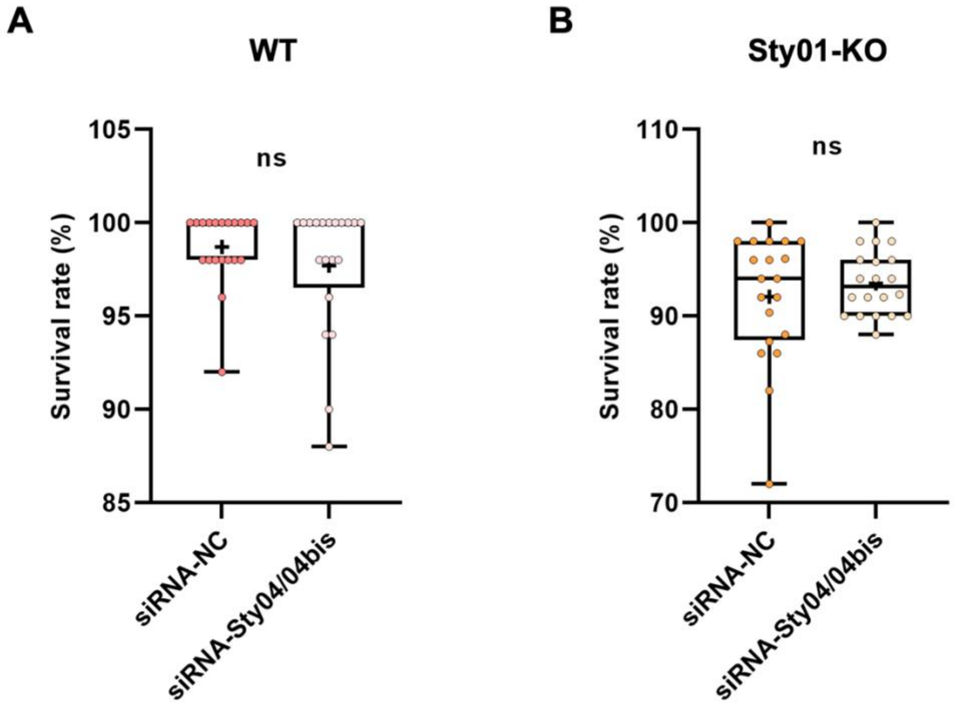


**Fig S5. Effect of *Stylin-04/04bis* gene silencing on the survival of *A. pisum* WT and Sty01-KO mutants.** Survival of aphids after 48 h of feeding on artificial diets supplemented with gene-specific siRNAs. Cohorts of WT (A) and Sty01-KO mutant (B) aphids were fed on artificial diets supplemented with siRNAs targeting *Stylin-04/04bis* (siRNA-Sty04/04bis) or a negative control siRNA (siRNA-NC). Bars show the minimum and maximum values, (+) indicate the mean of 20 independent replicates. Statistical analysis was assessed using generalized linear mixed models (GLMMs); ns, not significant (*P* > 0.05).

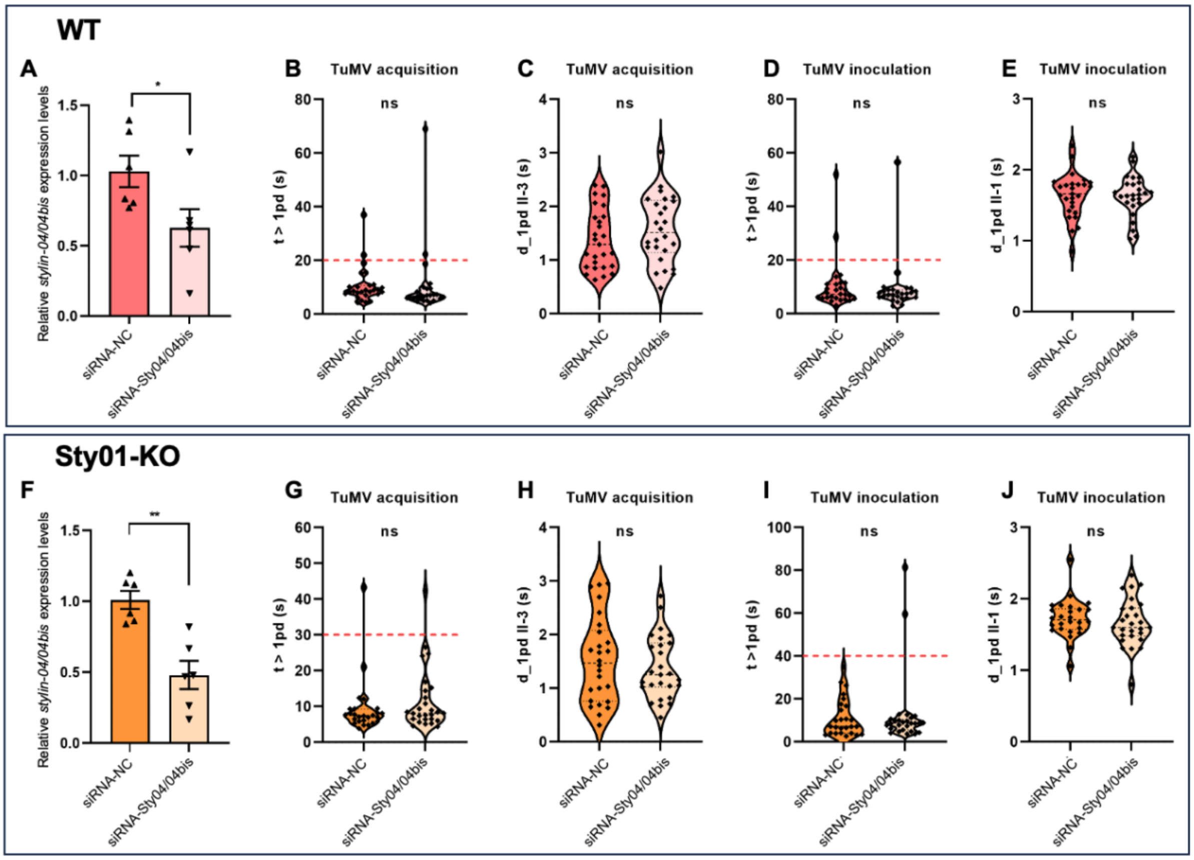


**Fig S6. Feeding behavior of wild-type and Sty01-KO mutant *A. pisum* lines during TuMV acquisition and inoculation after silencing *stylin-04/04bis*.**

Short-term feeding behavior of wild-type and mutant nymphs fed for 48 hours on artificial diets supplemented with specific siRNA targeting *Stylin04/04bis* expression (siRNA-Sty04/04bis) or negative control siRNA (siRNA-NC) was monitored by electrical penetration graph (EPG) analysis. (A-E) Data retrieved from WT treated aphids. (F-J) Data retrieved from Sty01-KO treated aphids. Relative *Stylin-04/04bis* expression levels of WT (A) and Sty01-KO aphids (F) exposed for 48 hours to siRNA artificial diets (mean ± SE; N = 6 pools of 3 aphids; asterisks show significant differences according to Student's *t*-test, $*P < 0.05$; $**P < 0.01$). First-instar nymphs were used at the start of the experiment. EPG variables associated with acquisition on infected turnips (B-C, G-H) or inoculation on healthy plants (D-E, I-J) were analyzed (see Table S3). Variables related to TuMV acquisition: (B, G) time from the beginning of the first probe to the first intracellular puncture (t > 1pd) and (C, H) duration of sub-phase II-3 within the first pd (d_1pd II-3) (N = 26-27, Mann-Whitney U test; ns, not significant). Variables related to TuMV inoculation: (D, I) t > 1pd and (E, J) duration of sub-phase II-1 within the first pd (d_1pd II-1) (N = 26-27, Mann-Whitney U test; ns = not significant). In panels (B, D, G, I), aphids plotted below the dotted line reached the first intracellular puncture within 30 s of recording onset. No significant difference was observed in the feeding behaviors associated with TuMV acquisition on infected plants or TuMV inoculation on healthy plants, regardless of the treatment and aphid lines.

**Table S1. Comparison of the feeding behavior of N2 nymphs of WT and mutant *A. pisum* lines.** Feeding behavior recorded for 10 min on turnip plants infected with TuMV (21 dpi) for EPG variables related to virus acquisition, and for 10 min on young healthy turnip plants for EPG variables related to virus inoculation. Time is expressed in seconds (mean ± SE).

| Variables | Aphid line | VIRUS ACQUISITION | | | VIRUS INOCULATION | | |
|---|---|---|---|---|---|---|---|
| | | N | WDI | *p* | N | WDI | *p* |
| **t > 1pd** | WT | 25 | 8.60±1.38 | 0.32 | 25 | 13.66±4.82 | 0.86 |
| | Sty01-KO | 25 | 7.69±0.90 | | 25 | 14.60±6.46 | |
| | Sty01-Cter | 22 | 5.98±0.29 | | 22 | 9.40±1.16 | |
| **d_1pd** | WT | 25 | 3.97±0.23 | 0.43 | 25 | 4.22±0.20 | 0.23 |
| | Sty01-KO | 25 | 4.31±0.14 | | 25 | 3.80±0.16 | |
| | Sty01-Cter | 22 | 4.02±0.16 | | 22 | 3.75±0.18 | |
| **d_1pd II-1** | WT | 25 | 1.27±0.07 | 0.99 | 25 | 1.35±0.05 | 0.12 |
| | Sty01-KO | 25 | 1.27±0.06 | | 25 | 1.32±0.05 | |
| | Sty01-Cter | 22 | 1.28±0.05 | | 22 | 1.42±0.06 | |
| **d_1pd II-2** | WT | 25 | 1.25±0.07 | 0.08 | 25 | 1.21±0.06 | 0.30 |
| | Sty01-KO | 25 | 1.39±0.07 | | 25 | 1.28±0.05 | |
| | Sty01-Cter | 22 | 1.18±0.07 | | 22 | 1.28±0.08 | |
| **d_1pd II-3** | WT | 25 | 1.45±0.16 | 0.48 | 25 | 1.66±0.15 | 0.01 |
| | Sty01-KO | 25 | 1.65±0.15 | | 25 | 1.20±0.11 | |
| | Sty01-Cter | 22 | 1.56±0.11 | | 22 | 1.05±0.09 | |

WDI: Waveform duration per insect.
t > 1pd: time from the beginning of the first probe to first intracellular puncture
d_1pd: duration of the first intracellular puncture.
d_1pd II-1, d_1pd II-2, d_1pd II-3: duration of subphases II-1, II-2 and II-3 indicated here for the first intracellular puncture.
*P*-values according to a Kruskal-Wallis test (*P* ≤0.05 considered statistically significant).

**Table S2. List of oligonucleotide siRNAs used in this study to silence *stylin* genes.**

| Gene targeted | Name | Type | Sequence (5' to 3') | Size |
|---|---|---|---|---|
| *Stylin-02* | siRNA-Sty02 | Sense | CCAAGAGUCUCUCAAAUUA (dTdT) | 21 |
| | | Antisense | UAAUUUGAGAGACUCUUGG (dTdT) | 21 |
| *Stylin-03* | siRNA-Sty03 | Sense | CACCAAUCAGCGUCAAAUA (dTdT) | 21 |
| | | Antisense | UAUUUGACGCUGAUUGGUG (dTdT) | 21 |
| *Stylin-04/04bis* | siRNA-Sty04/04bis | Sense | CAGCGUUUCUACAACAACG (dTdT) | 21 |
| | | Antisense | CGUUGUUGUAGAAACGCUG (dTdT) | 21 |

**Table S3. Comparison of the feeding behavior of N2 nymphs of WT and Sty01-KO lines after silencing *Stylin-04/04bis.*** Feeding behavior recorded for 10 min on turnip plants infected with TuMV (21 dpi) for EPG variables related to virus acquisition, and for 10 min on young healthy turnip plants for EPG variables related to virus inoculation. Time is expressed in seconds (mean ± SE).

| Variables | Aphid line | Treatment | VIRUS ACQUISITION | | | VIRUS INOCULATION | | |
|---|---|---|---|---|---|---|---|---|
| | | | N | WDI | *p* | N | WDI | *p* |
| **t > 1pd** | WT | siNC | 27 | 10.52±1.30 | 0.0503 | 27 | 10.31±1.86 | 0.68 |
| | | siSty04 | 26 | 10.27±2.49 | | 26 | 9.25±1.95 | |
| | Sty01-KO | siNC | 26 | 9.24±1.51 | 0.25 | 26 | 11.17±1.73 | 0.79 |
| | | siSty04 | 27 | 11.53±1.68 | | 27 | 12.65±3.30 | |
| **d_1pd** | WT | siNC | 27 | 4.00±0.15 | 0.31 | 27 | 3.93±0.17 | 0.78 |
| | | siSty04 | 26 | 4.20±0.14 | | 26 | 4.00±0.18 | |
| | Sty01-KO | siNC | 26 | 4.11±0.20 | 0.70 | 26 | 4.21±0.16 | 0.42 |
| | | siSty04 | 27 | 3.96±0.13 | | 27 | 3.96±0.18 | |
| **d_1pd II-1** | WT | siNC | 27 | 1.58±0.05 | 0.27 | 27 | 1.64±0.06 | 0.87 |
| | | siSty04 | 26 | 1.64±0.04 | | 26 | 1.63±0.06 | |
| | Sty01-KO | siNC | 26 | 1.61±0.04 | 0.77 | 26 | 1.72±0.05 | 0.38 |
| | | siSty04 | 27 | 1.63±0.06 | | 27 | 1.67±0.06 | |
| **d_1pd II-2** | WT | siNC | 27 | 1.03±0.05 | 0.40 | 27 | 1.01±0.06 | 0.84 |
| | | siSty04 | 26 | 0.99±0.04 | | 26 | 0.98±0.04 | |
| | Sty01-KO | siNC | 26 | 0.98±0.04 | 0.71 | 26 | 1.01±0.04 | 0.46 |
| | | siSty04 | 27 | 0.95±0.04 | | 27 | 1.06±0.06 | |
| **d_1pd II-3** | WT | siNC | 27 | 1.39±0.11 | 0.32 | 27 | 1.28±0.11 | 0.46 |
| | | siSty04 | 26 | 1.57±0.12 | | 26 | 1.39±0.11 | |
| | Sty01-KO | siNC | 26 | 1.52±0.16 | 0.74 | 26 | 1.48±0.13 | 0.13 |
| | | siSty04 | 27 | 1.38±0.11 | | 27 | 1.23±0.12 | |

siNC: Aphids fed on artificial diets containing siRNA-NC (negative control) (final concentration of 1 µmol).
siSty04: Aphids fed on artificial diets containing siRNA-Stylin-04/04bis (final concentration of 1 µmol).
WDI: Waveform duration per insect.
t > 1pd: time from the beginning of the first probe to first intracellular puncture time from the beginning of
d_1pd: duration of the first intracellular puncture.
d_1pd II-1, d_1pd II-2, d_1pd II-3: duration of subphases II-1, II-2 and II-3 indicated here for the first intracellular puncture.
*P*-values according to a Mann-Whitney U test ($p \leq 0.05$ would be considered statistically significant).

**Table S4**. **List of the oligonucleotide primers used for qRT-PCR analyses.**

| Name | Gene ID | Sequence (5' to 3') | Amplicon size | Reference |
|---|---|---|---|---|
| **Mdh2-F**[a] | NM_001126203.2 | TCCTGTTATCGGCGGACATT | 176 | Fu et al., 2025 |
| **Mhd2-R** | | ATGCCATAGACAGGGTTGCA | | |
| **qSty02-F**[b] | NM_001162671.2 | GGTCACATTAGTCCTCTTCATCG | 107 | Deshoux et al., 2020 |
| **qSty02-R** | | TGACTTCTTGTTCTTGGGACAG | | |
| **qSty03-F**[c] | NM_001161959.2 | ATCGTCAGTCACACTCGAGAAG | 103 | Deshoux et al., 2020 |
| **qSty03-F** | | GAGTCGTGGGTTGAAACTGTTG | | |
| **qSty04-F**[d] | NM_001172260.1/ NM_001172268.2 | ATCGCCTCCGTCGCCTTATTTC | 135/159 | Fu et al., 2025 |
| **qSty04-R** | | GGGTTTTGCCCGTTGTTGTAG | | |

[a] *Mitochondrial malate dehydrogenase*
[b] *Stylin-02*
[c] *Stylin-03*
[d] *Stylin-04/04bis*


Fu Y, Deshoux M, Cayrol B, Le Blaye S, Achard E, Hudaverdian S, Cloteau R, Pichon E, Strozyk E, Prunier-Leterme N, Jousselin E, Sauvion N, Thébaud G, Le Trionnaire G, Colella S, Uzest M. 2025. Stylet cuticular gene-directed mutagenesis impairs the pea aphid vector capacity to transmit a plant virus. PLoS Pathog. 2025 May 23;21(5):e1013192. doi: 10.1371/journal.ppat.1013192.

Deshoux M, Masson V, Arafah K, Voisin S, Guschinskaya N, van Munster M, et al. 2020. Cuticular Structure Proteomics in the Pea Aphid *Acyrthosiphon pisum* Reveals New Plant Virus Receptor Candidates at the Tip of Maxillary Stylets. J Proteome Res. 2020/01/29 ed. 2020;19: 1319–1337. doi:10.1021/acs.jproteome.9b00851
